# Data mining of digitized health records in a resource-constrained setting reveals that timely immunophenotyping is associated with improved breast cancer outcomes

**DOI:** 10.1101/198721

**Authors:** Arturo López-Pineda, Mario F. Rodríguez-Moran, Cleto Álvarez-Aguilar, Sarah M. Fuentes Valle, Román Acosta-Rosales, Ami S. Bhatt, Shruti N. Sheth, Carlos D. Bustamante

## Abstract

**Background:** Organizations that issue guidance on breast cancer recommend the use of immunohistochemistry (IHC) for providing appropriate and precise care. However, little focus has been directed to the identification of maximum allowable turnaround times for IHC, which is necessary given the diversity of hospital settings in the world. Much less effort has been committed to the development of digital tools that allow hospital administrators to monitor service utilization histories of their patients.

**Methods:** In this retrospective cohort study, we reviewed electronic and paper medical records of all suspected breast cancer patients treated at one secondary-care hospital of the Mexican Institute of Social Security (IMSS), located in western Mexico. We then followed three years of medical history of those patients with IHC testing.

**Results:** In 2014, there were 402 breast cancer patients, of which 30 were tested for some IHC biomarker (ER, PR, HER2). The subtyping allowed doctors to adjust (56.7 %) or confirm (43.3 %) the initial therapeutic regimen. The average turnaround time was 56 days. Opportune IHC testing was found to be beneficial when it was available before or during the first rounds of chemotherapy.

**Conclusions:** The use of data mining tools applied to health record data revealed that there is an association between timely immunohistochemistry and improved outcomes in breast cancer patients. Based on this finding, inclusion of turnaround time in clinical guidelines is recommended. As much of the health data in the country becomes digitized, our visualization tools allow a digital dashboard of the hospital service utilization histories.

## 1. Background

In breast cancer, immunohistochemistry (IHC) is a critical part of the accepted standard of care for determining prognosis and response to therapy. A panel of three IHC markers—estrogen receptor (ER), progesterone receptor (PR), and human epidermal growth factor receptor 2 (HER2)—are the most commonly used due to its predictive value for chemotherapy response in breast cancer [1]. Despite the existence of other IHC markers (e.g. Ki67, p53)[2], and the development of several commercial multi-gene tests that improve prognosis and treatment selection[3], the availability and expense of these tests remain largely prohibitive for constrain-resourced hospital settings, regardless of country income.

Mexico is an emerging country with an upper-middle income economy (as considered by the World Bank), where the healthcare system still has challenges procuring enough resources. The Mexican consensus [4] and clinical practice guidelines for breast cancer [5-7], the main working documents for oncologists in the country, recommend routine use of a 3-marker panel (ER, PR, HER2) to guide management and treatment for patients. In the Mexican population, the estimated frequency of ER/PR positive tumors is 60%, HER2 positive tumors is 20%, and triple negative tumors is 23% [8,9]. While these frequencies mirror those of other high and middle income countries, there is a troubling onset of disease among younger patients in Mexico (aged <40), with a high prevalence of triple-negative breast cancers [10]. Furthermore, there is an increased mortality trend associated with breast cancer in the country [11]. Timely testing for these IHC biomarkers needs to be fully recognized in clinical guidelines and hospital policies, and the testing must account for the wide diversity of patient trajectories and hospital settings [12].

Recently, organizations that issue guidance on breast cancer care worldwide have been broadening their focus to include resource-stratified guidelines, which reflects the diversity of hospital settings in the world:

- The Breast Health Global Initiative (BHGI) guideline for breast cancer healthcare in low-income and middle-income countries [13] offers four levels of guidance: basic (hormone receptor status by empiric assessment or response to treatment); limited (determination of ER status by IHC); enhanced (determination of PR status by IHC and measurement of HER2 overexpression); and maximal (gene profiling testing).
- The National Comprehensive Cancer Network (NCCN) framework for oncology care resource-stratification [14] offers four levels of guidance equivalent to those in the BHGI guideline.
- The American Society for Clinical Oncology (ASCO) offers a resource-stratified guideline for cervical cancer [15], which can be used to illustrate the need for a similar guideline for breast cancer.
- The World Health Organization (WHO) [16] offers guidance to countries on three resource scenarios (low, medium, high) with differential actions in the national cancer control programs, but has not yet offered a breast cancer specific resource-stratified guideline.

In this study, we aimed to quantify the potential effect that timely IHC testing has in improving patient outcomes. We hypothesized that with the use of data mining on digitized hospital records it is possible to facilitate the quantification of IHC testing as well as other services offered to breast cancer patients. At the same time, this study provided hospital administrators with a visual monitoring tool that facilitates the burden of human-intensive labor to quantify critical procedures in resource-constrained hospitals (including IHC testing).

## 2. Methods

### Hospital setting

The Mexican Institute of Social Security is a hybrid single-payer system with an integrated network of hospitals nationwide. In Michoacan (a state located in western Mexico), the General Regional Hospital No. 1 (HGR1) is the designated secondary-care facility for the IMSS-insured population, estimated at 1,288,695 people (28% of the state’s population), according to the 2015 census [17]. While there is a lot of heterogeneity between states in Mexico, it is important to highlight that the state of Michoacan ranks 29 out of 32 states in terms of competitiveness, governmental efficiency, and overall wealth [18].

The HGR1 hospital is located in a suburban community adjacent to Morelia (the capital of Michoacan). HGR1 is the secondary-care facility that serves as reference for seven General Zone Hospitals and 45 Family Medicine Units within the state of Michoacan. HGR1 has an oncology unit with a full complement of fixed staff and facilities available to all patients. It also offers pathology services, diagnostic imaging, and therapeutic capabilities with access to all approved drugs. Patients with breast cancer who require chemotherapy are sent to the outpatient medical unit, and patients with breast cancer who require radiation therapy are provided the service through an outsourced private service in the same city. The personnel include gynecologists, medical oncologists, surgical oncologists with significant breast cancer training, adequate numbers of nursing and pharmacy staff, surgeons with significant training, and cancer pathologists.

### Patients

The study population for this research was all cumulative breast cancer patients seen in 2014 by medical staff at HGR1, as reported by the institution’s breast cancer census in the absence of a cancer registry. To avoid selection bias in our study, we did not restrict the selection of patients to members of the female sex or any age group.

The general process at IMSS for breast cancer diagnosis is as follows. A patient with suspected breast cancer is referred from a primary-care facility to HGR1 after being examined by his/her family physician. If the physician suspects the existence of a breast lesion, the patient undergoes an imaging assessment using the breast imaging-reporting and data system (BIRADS) score. Upon arriving at HGR1, the patient is either seen by the breast cancer clinic (patients with BIRADS score 0 and 3) for further imaging studies and/or fine-needle biopsy assessments, or directly referred to the medical oncology service (patients with BIRADS score 4, 5 and 6), at which point they undergo surgical biopsies for pathology analysis. The gold-standard diagnosis of breast cancer is then provided by a board-certified pathologist, who assigns one of the C50 codes (malignant neoplasms of breast) from the international classification of diseases version 10 (ICD 10).

The final inclusion criteria were those whose biopsies had been tested with IHC to detect ER, PR, HER2 antibodies; then. We reviewed the medical records (both electronic and paper) of all breast cancer patients at HGR1, selected only those with IHC testing, and followed their medical histories, ending with March 2017, making note of multiple medical visits to the breast cancer clinic, medical oncology service, and/or surgical oncology service. From their medical records, we extracted information about IHC testing (antibody ordered, date of ordering, date of results being obtained), chemotherapy and hormonal drugs administered, and radiation sessions and surgical procedures undertaken.

### Study design

This was a retrospective hospital-based cohort study, using medical records collected routinely as part of clinical care. The objective was to understand the therapeutic impact on breast cancer patients of the time taken to test immunohistochemistry biomarkers in a resource-constrained hospital in western Mexico. We built a digital monitoring tool to report the frequency of treatment selection or treatment adjustment frequencies depending on breast cancer subtyping and the turnaround times for immunohistochemistry biomarker testing. The STROBE checklist is provided in Supplementary Material 1. All analyses were performed in R version 3.3.2.

#### Patient Trajectory Monitoring

We explored the paper- and electronic-based medical records for all patients with IHC testing, from the patient’s first day at the hospital to the last follow-up pertaining to this study (occurring before March 2017). First, author MFRM manually reviewed the patient’s medical records and curated a set of possible events experienced by the patients of this study, annotating the time points for each event for each patient. Then, author ALP annotated those events with standardized coding from the Unified Medical Language System (UMLS). When disagreement occurred, both authors discussed the translation to assign a UMLS code to the medical records in Spanish. Finally, similar events were grouped in top-level categories. The UMLS codes, UMLS descriptions, Spanish description, and the top-level categories can be seen in Table 1.

**Table 1.**
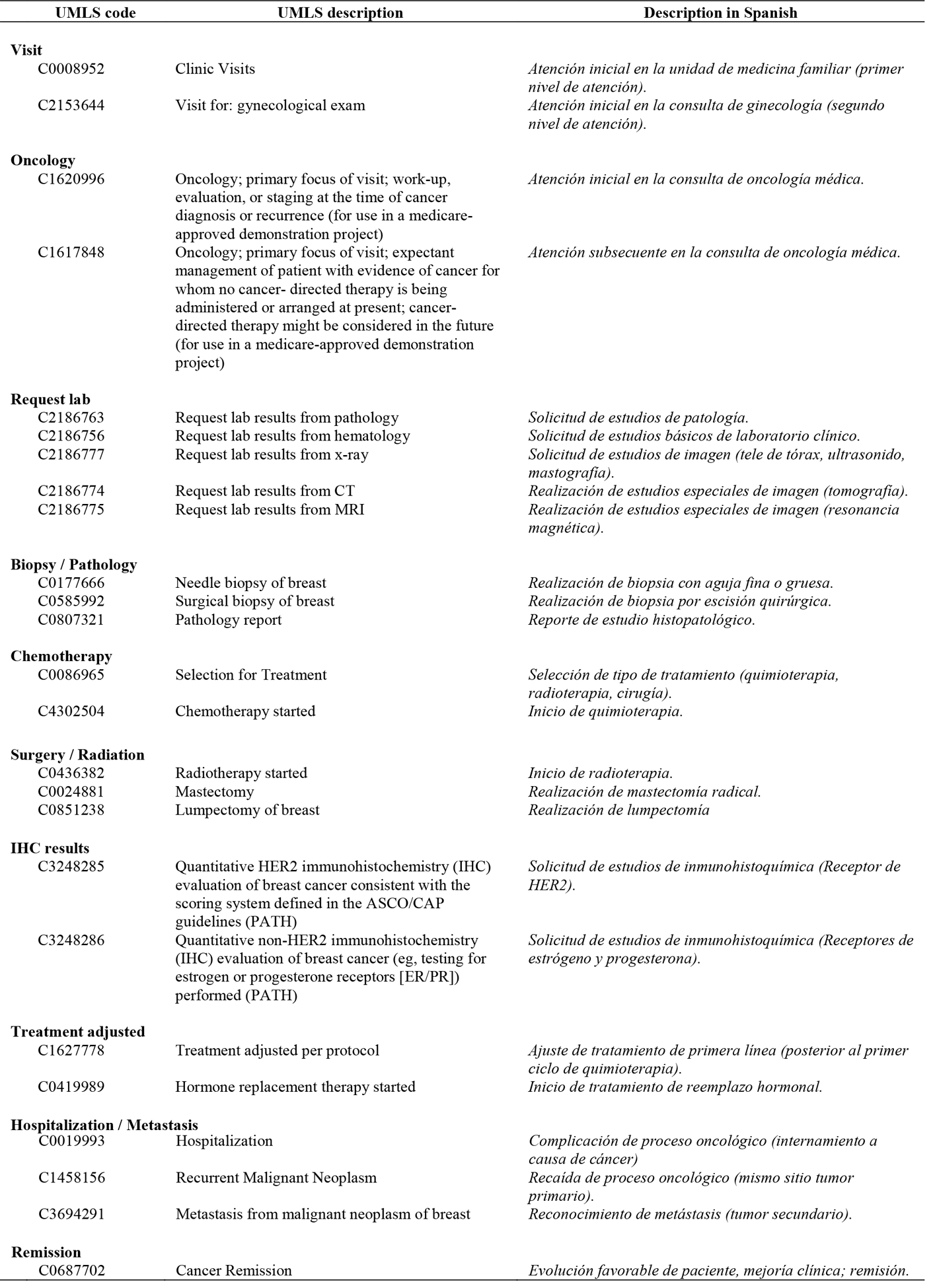
List of medical events with UMLS codes and descriptions in English and Spanish.

#### Missing data

Missing data was assumed to be non-missing at random (NMAR), and the events from Table 1 were considered missing for a specific reason. For example, if the code C2186775 (request lab result from MRI) was missing from a patient’s medical record, we left that value unassigned, since it was assumed that an MRI was not requested for that patient. There is a small chance that this assumption might be violated (and introduce some bias) if the physician did request the exam/procedure but forgot to write it down in the medical record. However, we did not impute any missing data, since that might be a larger source of bias for this EHR-based data.

#### Outliers

We assessed the time-to-event distribution for each event in Table 1, and reported the distribution of the patient cohort in violin plots that included: the median turnaround time, the interquartile probability density estimation, and the 95% confidence interval (whiskers). Outliers were identified using the 1.5 interquartile rule, but were not removed from the downstream analysis due to scarcity of data.

#### Clustering

A heatmap with all the patients and events was generated. We clustered patients using the k-means method, and added dendrograms to the heatmap. The distance matrix (needed to use this method) was generated with the time-to-event matrix, filling all missing events with a negative number (–1000). We tested for cluster stability to find the most appropriate value for k.

### 3. Results

Our data mining efforts revealed the epidemiological information pertaining to this IMSS hospital as shown in Table 2. In this cohort, all patients were female. The highest prevalence of cancer was within the 50 to 59 year-old age group. There were 207 patients (51%) diagnosed with early stages of cancer (stages I and II). Only 30 patients (7%) had undergone estimated subtyping with IHC testing. Of all patients, 18 patients (4%) were younger than 40 years old. At this hospital, 23 patients (6%) had a BIRADS score of 2 or lower, meaning they should not have been sent to the secondary care facility. In these cases, patients had external biopsy investigations in non-IMSS hospitals (usually private) and decided to continue their care for follow-up treatment at the IMSS oncology services.

**Table 2.** Clinical characteristics of the cohort and receptor status for patients with IHC testing

| Characteristic | Total (%) | With IHC testing (%) |
| --- | --- | --- |
| **Overall** | **N = 402** | **N = 30** |
| Sex |  |  |
| Female | 402 (100%) | 30 (100%) |
| Male | 0 | 0 |
| Age at diagnosis |  |  |
| <40 | 18 (4%) | 3 (10%) |
| 40-49 | 62 (15%) | 16 (53%) |
| 50-59 | 174 (43%) | 7 (23%) |
| 60-69 | 127 (32%) | 4 (13%) |
| 70+ | 21 (5%) | 0 |
| BIRADS |  |  |
| 0,1,2 | 23 (6%) | 18 (60%) |
| 3 | 203 (50%) | 7 (23.3%) |
| 4,5,6 | 162 (40%) | 4 (13.3%) |
| unknown | 14 (3%) | 1 (3.3%) |
| Tumor stage |  |  |
| I | 85 (21%) | 2 (7%) |
| II | 122 (30%) | 12 (40%) |
| III | 84 (21%) | 12 (40%) |
| IV | 19 (5%) | 3 (10%) |
| unknown | 92 (23%) | 1 (3%) |
| Patients without IHC testing | 372 (93%) | 0 |
| Patients with IHC testing | 30 (7%) | 30 (100%) |
| ER testing |  |  |
| positive | n.a. | 17 (57%) |
| negative |  | 11 (37%) |
| unknown |  | 2 (7%) |
| PR testing |  |  |
| positive | n.a. | 16 (53%) |
| negative |  | 12 (40%) |
| unknown |  | 2 (7%) |
| HER2 testing |  |  |
| positive | n.a. | 5 (17%) |
| negative |  | 23 (77%) |
| unknown |  | 2 (7%) |
| Additional IHC testing |  |  |
| Ki67, P53 | n.a. | 1 (3%) |
| unknown |  | 29 (97%) |
| Subtype according to the Mexican Consensus[4] |  | 14 (47%) |
| Luminal A | n.a. | 3 (10%) |
| Luminal B |  | 9 (30%) |
| Basal-like / Triple negative |  | 2 (7%) |
| HER2 |  | 2 (7%) |
| not enough information to assign |  |  |

### Digital Monitoring

Patient trajectories were finely segmented as shown in Figure 1, which further allow digital analysis (and therefore optimization), and clustered according to the groupings shown in Figure 2. Each patient is represented in a panel (rectangle) with colored bars, indicating the events that a patient experienced in the IMSS hospital. Each row represents one year of follow-up treatment for that patient, which can be from one to four rows (because the time maximum follow-up time span was three years and five months). The length of each colored bar represents the time between the occurrence of an event and the event that preceded it. Although events do not occur continuously, but, rather, happen at single time points (e.g. a patient visit, obtaining the results of a radiograph), the visual representation shown in Figure 1 provides a sense of how much time was required for each event to occur, assuming nothing else happened at the same time.

**Figure 1.**
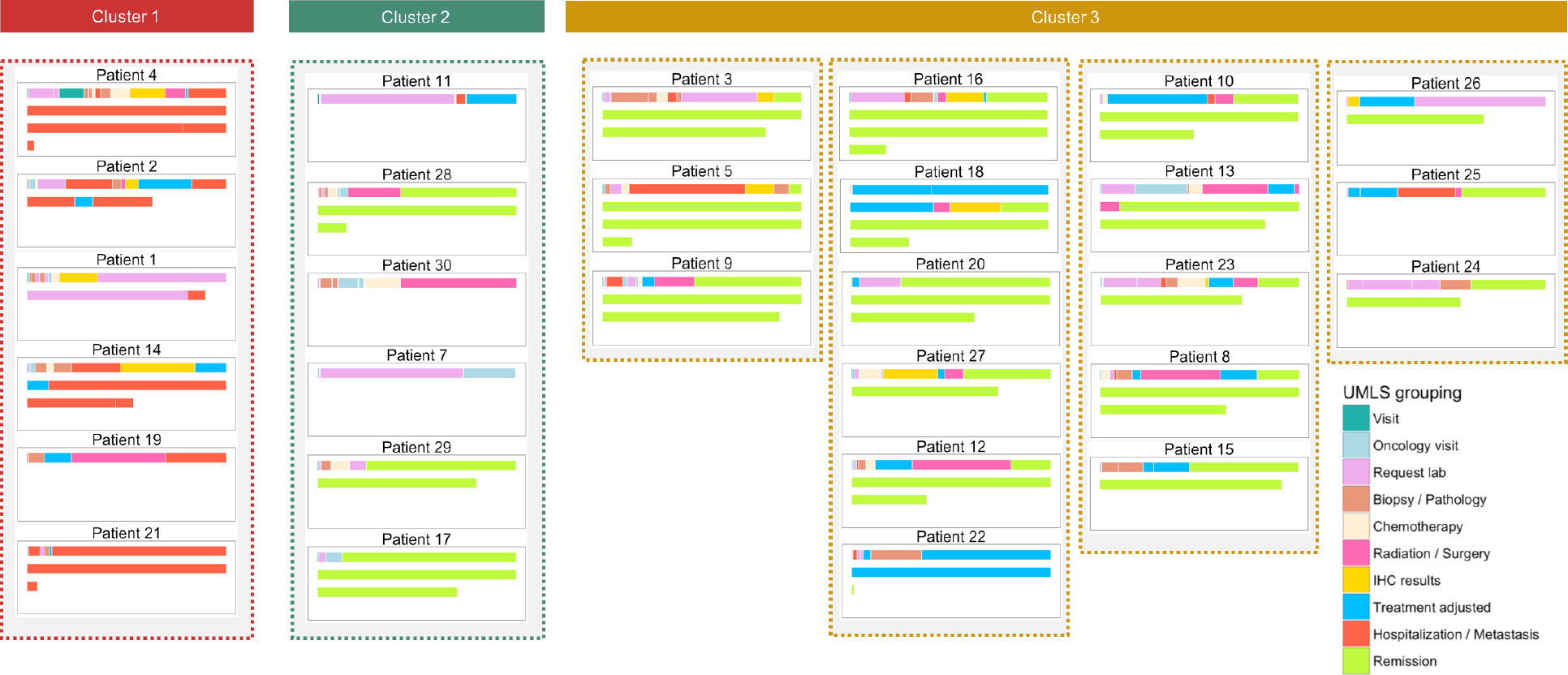
Medical trajectory plots of the cohort with IHC testing. Each patient is represented by a rectangle, with rows representing each year of follow-up treatment. Events are color-coded according to the type of event, and the length of the bar represents the duration between the occurrence of an event and the event that preceded it.

**Figure 2.**
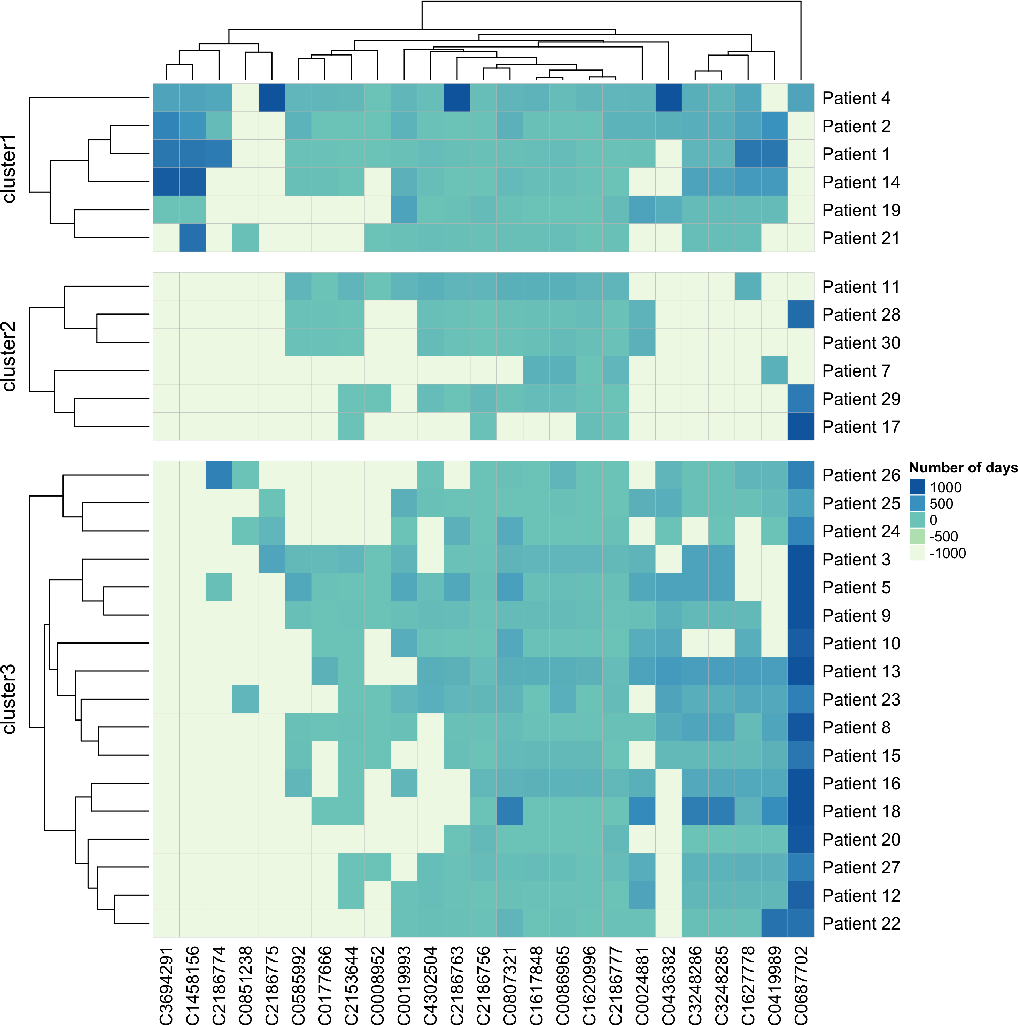
Heatmap of time-to-event with patient clustering. The time in which each event occurred for each patient is measured using information found in their medical record (in days, positive). To be able to create a dissimilarity matrix to calculate distance between patients, all missing values were assigned a negative value of -1000. Three main groups of patients (or clusters) were identified in this graph. This figure does not provide a context for the ordering in which events occurred, but it can still help to provide information about patients with similar trajectories.

### Turnaround time

Further data mining on this cohort identified the turnaround time for IHC testing as shown in Figure 3. Only 20 patients had information in their medical records relating to the timing of IHC testing. Although this service is referred to in the IMSS medical records, it is subcontracted to a private laboratory. There was large variation surrounding two key variables: a) the turnaround time to obtain results, with an average time of 56 days (95% C.I. 36 – 77 days); and b) the time between requesting IHC testing relative to the first day of follow-up treatment at the hospital, with an average time of 117 days (95% C.I. 80 – 154 days). Overall, for the 30 patients with IHC testing, 17 patients (56.7%) had their treatments adjusted, and 13 patients (43.3%) had their treatments confirmed.

**Figure 3.**
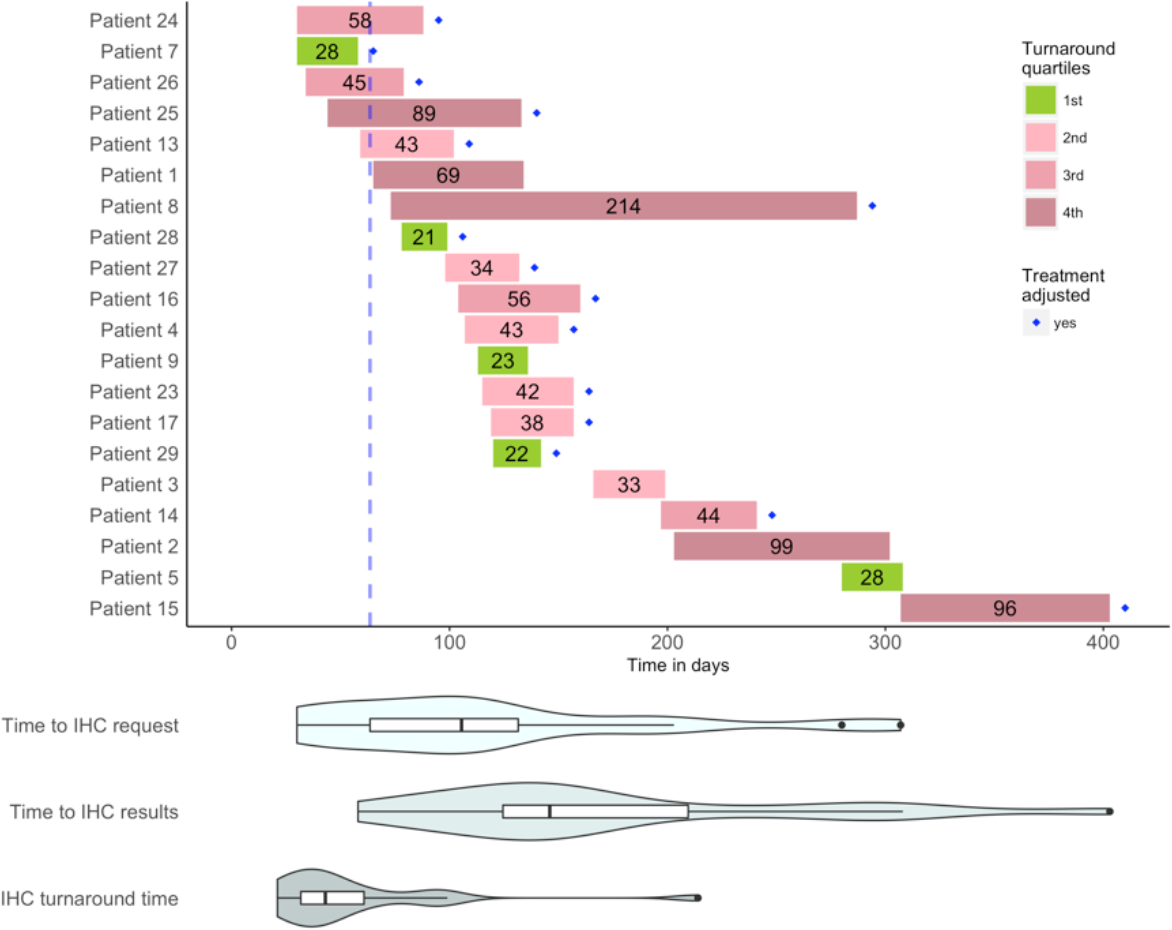
Turnaround times for IHC testing. The size of the bar indicates time in days from requesting results for IHC testing to obtaining them. For each patient, the initial time is the first day of follow-up treatment at the hospital. The color of the bar indicates the quartile within which the data falls in the distribution. The diamond mark indicates that the treatment was adjusted, according to protocols, after obtaining IHC results. The blue dashed line indicates the 1^st^ quartile for the time it took before an IHC request was submitted. Violin plots with whiskers (equivalent to a boxplot with a probability density estimation) provide an overall view of the turnaround time needed for IHC testing in three categories: a) from first day at hospital to request; b) from first day at hospital to obtaining IHC results; and c) from request of IHC to obtaining results.

## 4. Discussion

In Mexico, breast cancer continues to be a public health concern, even though more and more research is being done on its genomic changes and characteristics. Although our study investigated a small cohort of patients, the prevalence and epidemiological characteristics among our cohort are similar to what has previously been estimated for the country (in regards to age group and staging) [11]. Our study further investigates the use of digital health mining tools for evaluating the timeliness of IHC testing, so our conclusions can then be used to inform hospital administrators and public health officials around the world about the importance of deploying digital tools for assessing timely IHC testing.

### On the therapeutic value

The therapeutic importance of immunophenotyping (the use of IHC testing for subtyping of cancer patients) needs to be carefully considered and addressed in treatment plans by oncologists. The traditional (FEC) chemotherapy regimen recommended by the IMSS protocol, composed by 5-fluorouracil, epirubicin, and cyclophosphamide, could be adjusted according to the IHC biomarker testing.

Patients with estrogen positive (ER+) and/or progesterone positive (PR+) could benefit from anti-hormone or endocrine therapy to prevent recurrence, either aromatase inhibitors to block estrogen production (e.g. anastrozole, letrozole, or exemestane), or the selective estrogen receptor modulators (SERMs), which interfere with the ability of estrogen to stimulate the growth of breast cells (e.g. tamoxifene or toremifene).

Additionally, patients with HER2 positive (HER2+), in which tumors tend to grow and spread more aggressively, could benefit from a targeted therapy that could block HER2 (e.g. trastuzumab or pertuzumab). The use of in situ hybridization (ISH), instead of IHC, can be used to determine HER2 status with an overall concordance with IHC, and it may be more beneficial to use both [19]. Furthermore, HER2 status has been successfully incorporated into medical practice to guide treatment decisions for breast cancer patients. In fact, the American Society of Clinical Oncology/College of American Pathologists (ASCO/CAP) updated their 2013 guidelines to designate more patients as eligible for trastuzumab therapy, in accordance with ISH and IHC testing [20]. Also, the BHGI resource stratified guidelines recommend the use of trastuzumab at the enhanced level due to the high cost of the drug, as well as for the availability for testing. The IMSS’s medications chart [21] reports trastuzumab as an available drug for any HER+ positive patient in the country.

### On the timeliness of testing

Opportune IHC testing, understood as the availability of IHC results before the beginning of any treatment, is of critical importance. For the IMSS hospital in this study, IHC results were typically not timely opportune, but whenever they were available they triggered a response from the clinical oncology team to adjust patient treatments. The wide dispersion of time to request and turnaround time demonstrates the lack of standardization of this process. Many factors could have contributed to the delay in IHC testing, including: patient non-adherence to appointments; hospital overflow, resulting in delayed appointments; logistical problems between IMSS and the external laboratory service; and administrative problems within the hospital resulting in inadequate tracking of tests and results.

### On the need for improved clinical guidelines

It is important to establish an appropriate timeline for opportune IHC testing. For example, In the United States, the College of American Pathologists’ (CAP) guideline on turnaround time for biopsy testing is around two days for 90% of cases [22]. In fact, the joint guideline on HER2 testing from the American Society of Clinical Oncology (ASCO) and CAP recommends informing the patients about the expected turnaround time [23]. In Australia, a study revealed that the average turnaround time was between 4-5 days [24]. In Saudi Arabia, a study showed that 24% of cases fall outside the recommended CAP turnaround time [25]. The European Society for Medical Oncology (ESMO) published a survey on 24 European countries were the turnaround time was 10 days or less for 89% of laboratories [26]. In the United Kingdom, the Royal College of Pathologists recommend the use of key performance indicators to have a histopathology diagnosis within seven days of biopsy in 90% of cases [27].

Given their resource-available settings, clinical guidelines in developed countries do not have a recommendation regarding the maximum turnaround time that could be effective in a patient’s treatment trajectory. More importantly, they do not address the time between initial diagnosis, molecular testing, and the selection of treatment. In Mexico, clinical guidelines for breast cancer recommend testing for ER, PR, and HER2 as part of the histopathology study, but fail to provide guidance regarding turnaround time. IMSS maintains a medical procedures manual, which includes ten key indicators for breast cancer screening, diagnosis, and treatment [28]. This manual measures time-to-diagnosis in a 30-day period, including imaging through mammography and results of histopathology report. In addition, the institution also measures time-to-treatment measured from the date of diagnosis, which should be achieved within a 21-day period. Recognizing the resource constraints of IMSS, the implementation of a key indicator policy related to IHC testing might significantly reduce turnaround times.

### On the need for accurate electronic health records (EHR)

Missing information was common across the paper and electronic records in our study. Clinicians and administrative staff at the IMSS hospital are still getting used to the novel implementation of the EHR, which had an impact on our ability to better characterize this cohort. In the absence of a cancer registry, our best estimate of the disease is the institutional census. As the hospitals in Mexico, and elsewhere, continue to become more and more electronic, there is a need to develop better software tools to analyze the information obtained, including medical natural language processing and machine learning applications.

### On digital monitoring and visual representation

The monitor shown in Figure 1 quickly provides a patient history overview of hospital care. With more work on the user interface, we envision this tool could eventually represent a valuable visual aid in which a patient and their companions might be able to better communicate with their clinicians about the management of their disease. Currently, some of the challenges of this tool include the missing identification of events that are overlapping, and the impossibility of further elaborating on the details of each event. For a hospital administrator, Figure 1 can provide a quick overview of the patients seen in their hospital, which could be used as a decision-making tool, with the proper validation. This visual aid monitoring tool can be obtained from the repository at https://github.com/bustamante-lab/patientTrajectory

## 5. Conclusions

Adverse events in the trajectory of an oncological patient, which might include hospitalization related to cancer, recurrence of tumor, or metastasis, are extremely costly for the healthcare system. The use of IHC testing was shown in our study to help with the selection of precise treatment for patients (either by adjusting the treatment or confirming it). The time in which IHC testing is performed is of critical importance if we want to influence improved prognosis.

We have shown that the use of opportune immunohistochemistry testing is associated with beneficial therapeutic effect on breast cancer patients. The aims of any healthcare system should be the identification of earlier events that can have an impact on downstream events in the trajectory of an oncological patient’s treatment. In resource-constrained settings, it is important not only to consider alternatives to more costly diagnostics (e.g. genomic testing), but also to incorporate the regulatory and logistical aspects of implementing these tests.

IMSS must face the important challenge of continuing to improve their turnaround times, which should have a positive impact on the prognosis of their patients. As the Mexican healthcare system continues to transition from reactive to preventative care, the need for more IHC testing in breast cancer and other diseases will certainly allow for the further development of digital patient monitoring.

## Declarations

### Ethics approval

This research was reviewed and approved by the Mexican Institute of Social Security (IMSS)’s National Scientific Research Committee and its National Bioethics Committee, under protocol number R-2017-785-010, and locally by IMSS’ Local Research and Ethics Committee, under specific protocol number R-2017-1602-14 HGR1 Charo, Michoacan. Stanford’s Institutional Review Board provided a non-human subject determination under eProtocol: 40704. Researchers only received access to non-identifiable data under the considerations of the applicable Mexican bylaws on health research, patient privacy, and electronic medical records. Consent was not required.

### Availability of data

Data belongs to the Mexican Institute of Social Security (IMSS), through its state Delegation of Michoacan. IMSS’s Coordination of Health Research may grant access to this data on a case-by-case basis to researchers who obtain the necessary approvals from IMSS’s National Scientific Research Committee and National Bioethics Committee.

### Competing interests

RAR declares to be IMSS’s state delegate in Michoacan, overseeing the hospital mentioned in this study. The remaining authors declare no conflicts of interest.

### Funding

Research reported in this publication was supported by grants U01 HG007436-04 from the National Human Genome Research Institute (NHGRI)’s Clinically Relevant Genome Variation Database; and U01 FD004979 from the Food and Drug Administration (FDA)’s joint Center of Excellence in Regulatory Science and Innovation (CERSI) at the University of California, San Francisco and Stanford University. The funders had no role in study design, data collection and analysis, decision to publish, or preparation of the manuscript.

### Authors’ contributions

ALP designed the study. MFRM reviewed medical notes. MFRM and ALP performed the analysis of the data. ASB, SNS, RAR, CAA, SMFV and CDB provided interpretation of the results. ALP drafted the manuscript, and all authors contributed critically, read, revised and approved the final version.

## Acknowledgements

The authors wish to acknowledge Katie Kanagawa, PhD, for her valuable support in editing this manuscript.

